# CpGeneAge: multi-omics aging clocks associated with Nf-κB signaling pathway in aging

**DOI:** 10.1101/2025.06.26.661785

**Authors:** Brigitta Varga, Csaba Kerepesi

**Author notes:** Correspondence: Csaba Kerepesi.

## Abstract

Aging clocks have emerged as the primary tools for measuring biological aging and have been developed for a wide range of single-omic measurements. Epigenetic aging clocks showed high accuracy in age prediction, however, their biological interpretation is still a challenging task. Transcriptomics aging clocks provide better interpretability but worse age prediction accuracy. To exploit the benefits of both omics techniques, the main goal of this study was to develop the first multi-omics aging clocks based on combined epigenetics and transcriptomics features. For this purpose, we utilized a dataset where reduced representation bisulfite sequencing (RRBS) and RNA-seq measurements were measured at the same time for peripheral blood samples of 182 individuals. Then we trained machine learning models (ElasticNet) using the methylation and gene expression features at the same time. While the most accurate models tended to use exclusively methylation features, we were able to develop highly accurate multi-omics aging clocks too (called CpGenAge). Both the canonical and the non-canonical Nf-κB signaling pathways, with the genes EDA, EDA2R, EDARADD, and CD70, were overrepresented among the gene expression features of the multi-omics aging clocks. The EDARADD, which is a unique hallmark of aging, was represented among both the gene expression and methylation features. By developing single-omic clocks on the same multi-omics dataset, we found that epigenetic age acceleration and transcriptomics age acceleration do not correlate with each other, further supporting the benefits of our multi-omics approach. In summary, here, we demonstrate that multi-omics aging clocks are useful tools to investigate aging and biological age at the multi-omics level.

## Introduction

Aging has a major impact on the economy and health, but its internal mechanisms are poorly understood^1,2^. Aging clocks (i.e., machine learning models that can predict age) have emerged as the primary tools for measuring biological age and have been developed for a wide range of omics measurements^3–5^. DNA methylation has proven to be the most effective omics measurement to develop biological aging clocks due to its predictive power. Despite the development of first, second, and third-generation epigenetic aging clocks for humans and model organisms based on microarrays^6–15^ and bisulfite-sequencing^16–23^, there is still room for improvements, and the biological interpretability of epigenetic clocks is still challenging^24^. Transcriptomics (i.e. gene expression-based) aging clocks also emerged for bulk^25–29^ and single-cell samples^30–33^ providing a better interpretability but worse age prediction accuracy compared to the methylation aging clocks. While aging clocks are typically trained on single-omic measurements, we hypothesized that using two different omics may provide a more accurate and interpretable approach for measuring biological age. Therefore, we developed multi-omics aging clocks using both the methylation and gene expression levels of each individual to capture epigenetics and transcriptomics signatures of aging at the same time.

## Results

### Development of multi-omics aging clocks

Our goal was to create accurate age prediction models with different omics biomarkers and investigate their role in aging. To achieve this, we used 182 peripheral blood samples with both reduced representation bisulfite sequencing (RRBS) and RNA-seq measurements (Bhak et al. dataset)^34^, and applying leave-one-out-cross-validation, we trained multi-omics aging clocks on a merged dataset containing expression of ∼21.600 genes and methylation levels of ∼4 million CpG sites (Fig. 1a). Increasing the strength of the lasso term of GLMNet the models tended to eliminate all transcriptomics features and used only DNA methylation features suggesting that methylation features have stronger predicting power than transcriptomics features. Models with alpha = 0 contained all features while models with alpha = 0.5 and 1 contained only methylation features with no gene expression features. We decided to select the parameter settings alpha = 0.01 as it resulted in models with some gene expression features besides the methylation features and it still provided accurate age predictions with mean absolute error (MAE) = 2.761 years and Pearson correlation coefficient (R) = 0.948 (Fig. 1b).

**Fig. 1.**
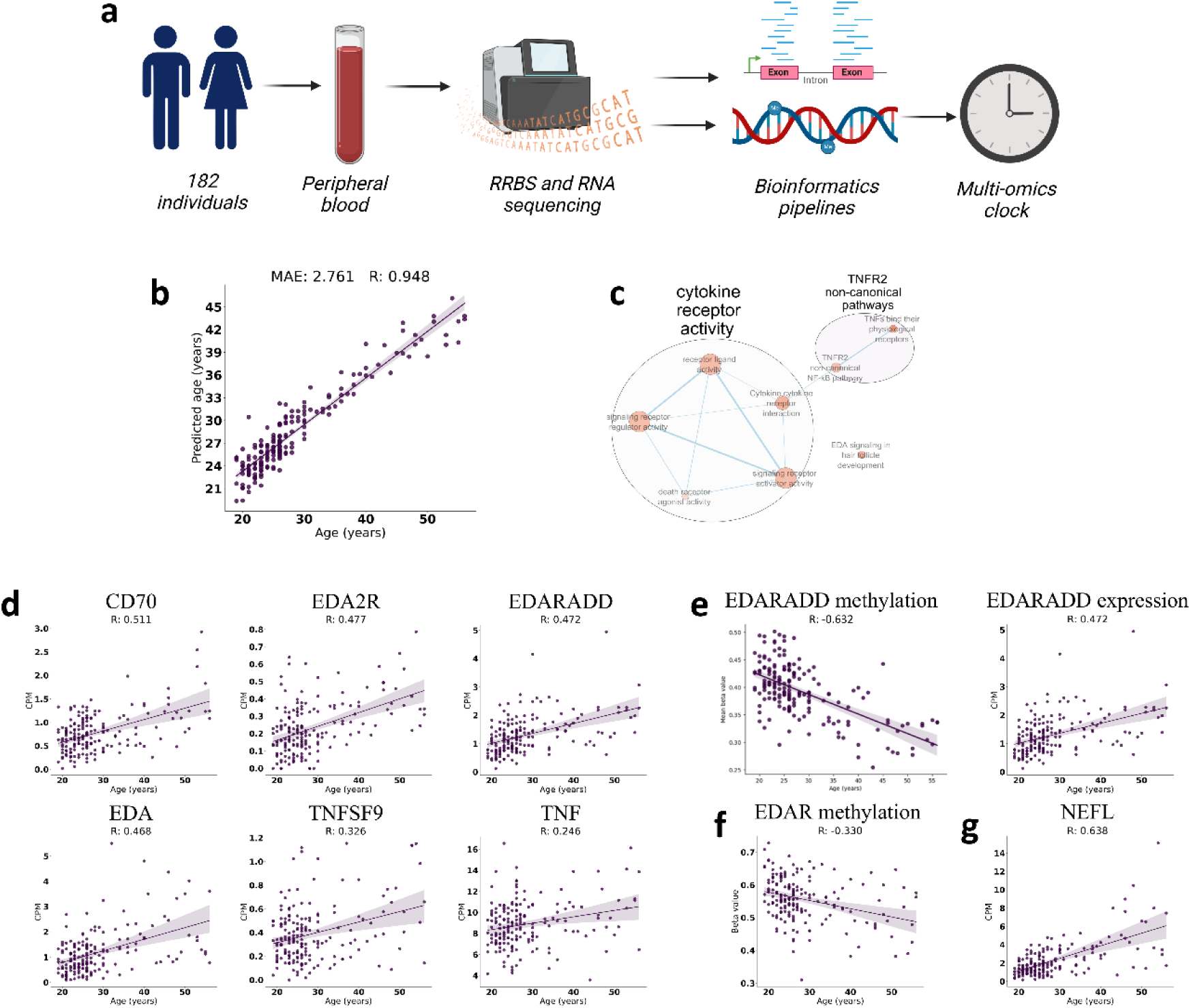
Development of multi-omics aging clocks based on CpG methylation and gene expression levels. **a,** Schematics of the multi-omics clock development. We processed RRBS and RNA-seq data of 182 peripheral blood samples (Bhak et al. dataset) and then developed multi-omics aging clocks. The figure was created with BioRender.com **b,** Performance of the multi-omics aging clocks by using leave-one-out cross-validation. **c,** Relationships of the most significantly overrepresented pathways and GO terms in the gene expression features (i.e. genes) of multi-omics aging clocks. The opacity of the nodes indicates significance of the overrepresentation (FDR-adjusted p-value), while the thickness of the edges corresponds to the number of common genes that we captured between the terms/pathways. The size of the circles represents the number of genes that belong to the terms/pathways on the labels. **d,** Association of chronological age and gene expression (measured in “counts per million”, CPM) for tumor necrosis factor receptors and multi-omics aging clock genes that belong to the Nf-κB pathways. **e,** The EDARADD gene is hypomethylated and upregulated in aging. **f,** CpG site of EDAR is hypomethylated in aging **g,** Neurofilament light chain gene expression had the highest correlation with age.

### Nf-κB signaling pathways were overrepresented among the gene expression features of the multi-omics aging clocks

To interpret biological functions captured by the multi-omics aging clocks we investigated their features. Besides the methylation features, 32 annotated gene expression features have been used by the 182 models of the leave-one-out-cross-validation and there were 18-23 genes in each model with high overlap among them (Table 1). We conducted an over-representation analysis (ORA) to find functions, processes, and pathways, based on the GO molecular functions, Reactome, WikiPathways, and KEGG databases, that are significantly enriched in the set of that 32 genes (Supplementary Table 1, and Fig. 1c). The most significantly enriched terms were “TNFs bind their physiological receptors” and “TNFR2 non-canonical NF-kB pathway” from the Reactome database; “Cytokine cytokine receptor interaction” and “EDA signaling in hair follicle development” from the WikiPathways; “Cytokine-cytokine receptor interaction” and “NF-kappa B signaling pathway” from the KEGG database; “signaling receptor regulator activity” and “receptor ligand activity” from the GO biological functions. These overrepresentations were caused predominantly by the presence of the genes CD70, EDA to EDA2R, and EDARADD, all of them associated with the Nf-κB pathways. These genes and the tumor necrosis factors were positively correlated with age (Fig. 1d).

**Table 1.** Gene expression features (i.e., genes) chosen by the multi-omics aging clocks.

| Gene symbol | Description | Pearson correlation gene expression and age |
| --- | --- | --- |
| NEFL | neurofilament light chain | 0.638 |
| LINC00824 | long intergenic non-protein coding RNA 824 | 0.596 |
| PI16 | peptidase inhibitor 16 | 0.574 |
| IGFBP3 | insulin like growth factor binding protein 3 | 0.565 |
| CRIP2 | cysteine rich protein 2 | 0.550 |
| HNRNPA1P21 | heterogeneous nuclear ribonucleoprotein A1 pseudogene 21 | 0.541 |
| ISM1 | isthmin 1 | 0.523 |
| CD70 | CD70 molecule | 0.511 |
| ADAM23 | ADAM metallopeptidase domain 23 | 0.505 |
| GREM2 | gremlin 2, DAN family BMP antagonist | 0.501 |
| PHLDA3 | pleckstrin homology like domain family A member 3 | 0.496 |
| PHF2P2 | PHD finger protein 2 pseudogene 2 | 0.493 |
| CHN1 | chimerin 1 | 0.485 |
| TPO | thyroid peroxidase | 0.478 |
| WNT10A | Wnt family member 10A | 0.478 |
| EDA2R | ectodysplasin A2 receptor | 0.477 |
| EDARADD | EDAR associated via death domain | 0.472 |
| CCR8 | C-C motif chemokine receptor 8 | 0.470 |
| TCEAL2 | transcription elongation factor A like 2 | 0.469 |
| EDA | ectodysplasin A | 0.468 |
| CAVIN1 | caveolae associated protein 1 | 0.425 |
| CACNA1I | calcium voltage-gated channel subunit alpha1 | 0.423 |
| LPAR3 | lysophosphatidic acid receptor 3 | 0.389 |
| TRHDE-AS1 | TRHDE antisense RNA 1 | 0.364 |
| PTPRH | protein tyrosine phosphatase receptor type H | 0.316 |
| LDHD | lactate dehydrogenase D | 0.298 |
| ROBO1 | roundabout guidance receptor 1 | -0.223 |
| TRBV6-4 | T cell receptor beta variable 6-4 | -0.284 |
| CACHD1 | cache domain containing 1 | -0.296 |
| NRCAM | neuronal cell adhesion molecule | -0.311 |
| CASC15 | cancer susceptibility 15 | -0.338 |
| LAMP5 | lysosomal associated membrane protein family member 5 | -0.355 |

The multi-omics aging clocks used 4150 CpG sites in total (Supplementary Table 5). The EDARADD gene covered 11 CpG site feature of the clocks and it was the only Nf-κB pathways gene that contained CpG sites in the methylation feature set of the multi-omics aging clocks. The mean methylation level of the 11 clock CpG sites on the EDARADD negatively correlated with age (Fig. 1e). The EDAR, which is the interaction partner of the EDARADD gene, was expressed but not correlated with age, however multi-omics clocks captured one demethylated CpG site of it. (Fig. 1f). Neurofilament light chain had the highest correlation with chronological age, which is a protein encoded by the NEFL gene (Fig. 1g).

### Comparison of the single-omics transcriptomics aging clocks and the multi-omics aging clocks

Investigating the overlap between the feature set of the multi-omics aging clock and single-omics transcriptomics aging clocks, we trained age prediction models using exclusively the gene expression features of the Bhak et al. dataset. In total 172 different genes were used by the 182 transcriptomics aging clock models (52-120 genes per model) with an accuracy of MAE = 3.98, and R = 0.826. Twenty-six gene expression features of the transcriptomics aging clocks overlapped with the 32 genes of the multi-omics aging clocks (Supplementary Table 3). Twenty of the 26 overlapped genes were among the top 21 highest age-correlated genes of the transcriptomics aging clocks. Interestingly, genes with a negative correlation with age were not present in the transcriptomics models, however, the multi-omics models contained such genes. Overrepresentation analysis had been conducted by using the 149 annotated genes of 172 gene expression features of the transcriptomics aging clocks. The Nf-κB pathways remained significantly overrepresented in this wider gene set too, however, we observed more diverse pathways enriched in the gene set of the transcriptomics aging clocks (Supplementary Table 4). In summary, multi-omics aging clocks tend to choose genes that have the highest correlation with age, while the transcriptomics clocks use an extended set of genes including ones with low correlations with age.

### Genes of the whole blood-based multi-omics aging clocks correlated with age in external PBMC datasets

To further investigate the gene expression features of the multi-omics aging clocks, we used an external validation dataset of healthy individuals. The Asian Immune Diversity Atlas (AIDA) dataset (v2) contained 1,265,624 immune cells, and 32 cell types from 625 healthy donors with ages between 19 and 75 years^35^. We constructed pseudobulk samples from the AIDA dataset by aggregating different cell-type-specific raw read counts per individual. We found that out of the features of the multi-omics aging clocks IGFBP3, PHILDA3, and NEFL had the highest positive correlation with age and further 26 genes correlated with age (Fig. 2a). We tested the previously trained whole blood-based single-omics transcriptomics aging clock on the AIDA pseudobulk PBMCs dataset for external validation and observed good generalization (Fig. 2b). Twenty-five features of the 52 transcriptomics aging clock features overlapped with the 32 genes of the multi-omics aging clock in the AIDA dataset including EDA, EDA2R, and EDARADD (Supplementary Table 2, and Fig. 2c). We used another validation dataset (Morandini et al. dataset)^36^ containing 143 high-quality samples of bulk peripheral blood mononuclear cells (PBMCs) from healthy individuals between ages of 20-74 years. We found that 14 of the 32 multi-omics aging clock genes correlated with age including EDA and EDARADD (Supplementary Table 2, and Fig. 2de), however, their correlations were lower than in the training set (Bhak et al. dataset) and the AIDA pseudobulk dataset.

**Fig. 2.**
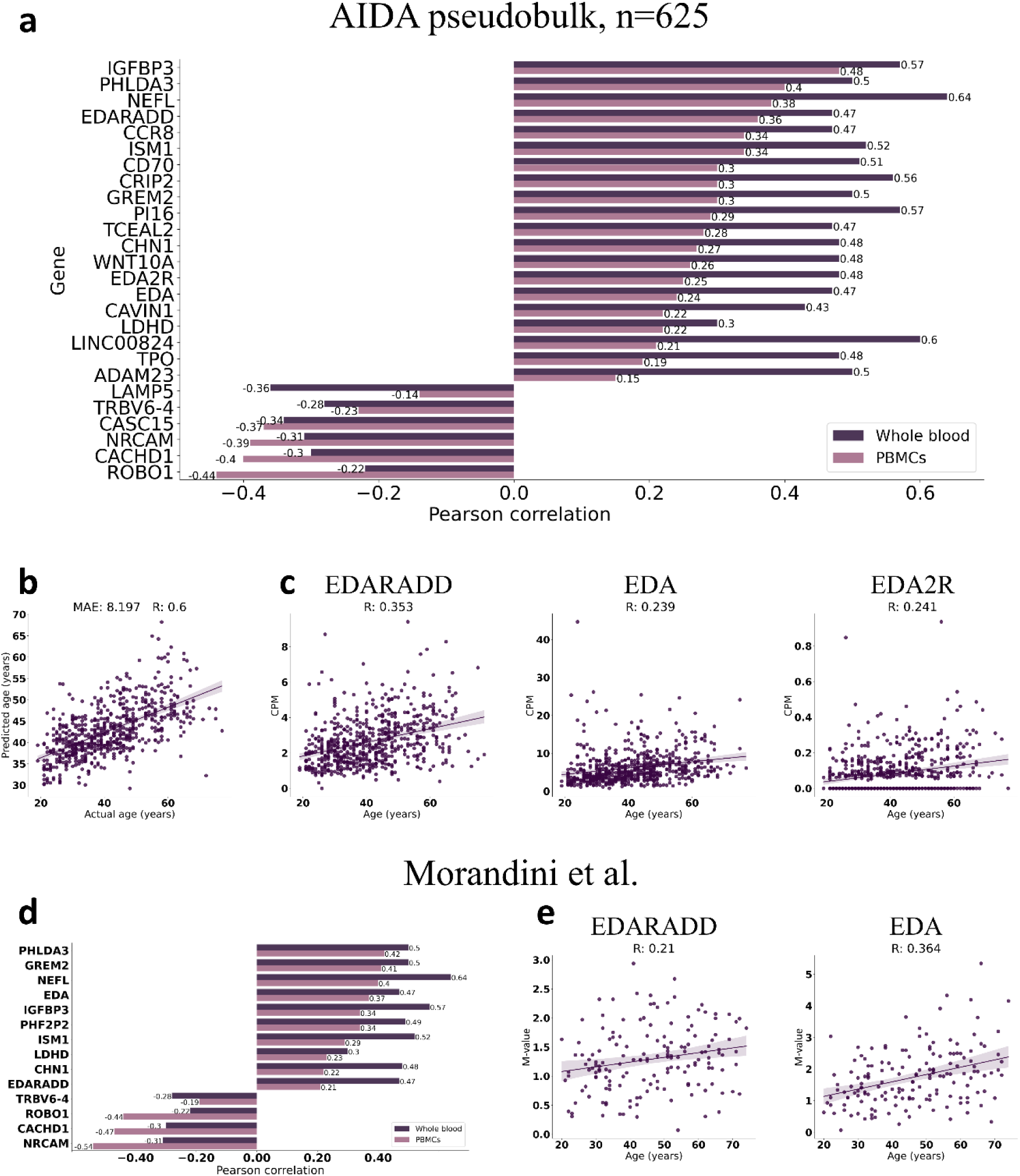
Genes of the whole blood-based multi-omics aging clocks correlated with age in external PBMC datasets. **a,** Correlation of chronological age and expression of the multi-omics aging clock genes in pseudobulk PBMC samples of the external dataset, AIDA. We also displayed the correlations in the whole blood bulk Bhak et al. dataset. **b,** The whole-blood-based single-omic transcriptomics clock performance tested on the pseudobulk PBMC samples of the external dataset, AIDA. **c,** Correlation of age and the gene expression of EDA, EDARADD, EDA2R in AIDA. **d,** Correlation of age and expression of the multi-omics clock genes calculated using PBMC samples of the Morandini et al. external dataset. We also displayed the correlations in the Bhak et al dataset (bulk whole blood). **e,** Correlation of age with gene expression of EDA, and EDARADD in the Morandini et al. PBMCs dataset.

### Promoter analysis of EDARADD

As we showed above, the EDARADD gene were shared between the two omics features of the multi-omics aging clock, and gene expression of EDARADD positively correlated with age while the mean methylation value of the 11 clock CpG sites of EDARADD negatively correlated with age (Fig 1e). We also investigated the 11 CpG sites individually and found that all of the CpG sites negatively correlated with age (Fig. 3b). Using a transcript from the MANE Project^37^, matched NCBI and EMBL-EBI transcripts and, we found three promoter CpG sites, which are located upstream -3, -62, -74 nt from the first nucleotide of the first exon., and the rest in the first intron (Fig. 3a). For further investigation, using a transcript from Havana/Ensemble automatic annotation merge, we found 6 promoter CpG sites that are located within -1500 nt upstream from from the first nucleotide of the first exon (TSS1500), -1259, -1247, -1188, -1055, -913, -787 and the rest 5 CpG sites are located downstream from the first exon, located in the first intronic region (Supplementary Table 6).

**Fig. 3.**
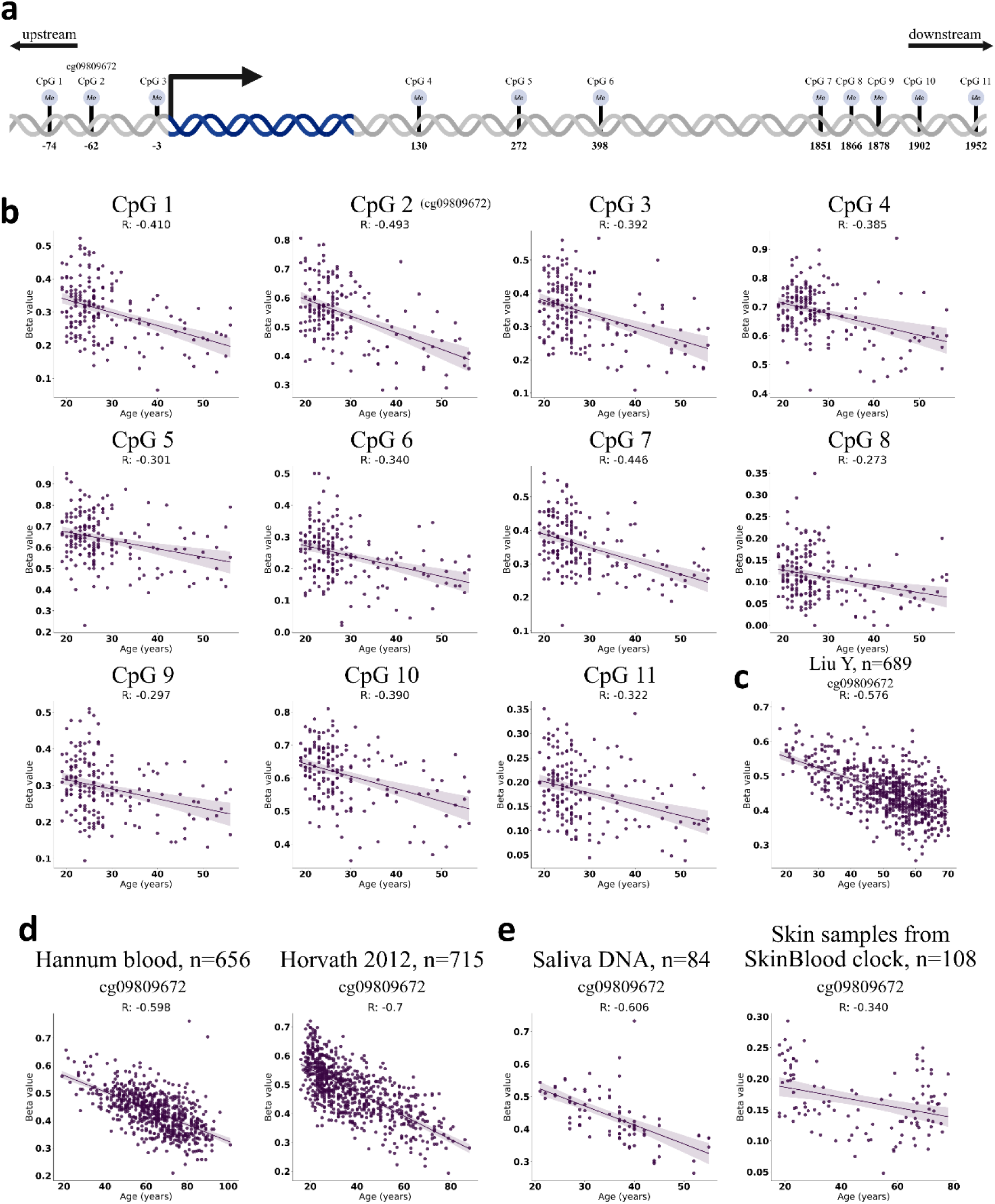
Promoter analysis of EDARADD. **a,** Schematics of EDARADD gene according to MANE selected consensus transcript. The arrow denotes the beginning of the coding region, the first nucleotide of the first exon and the direction of the transcription. Blue area denotes the coding region while the grey is the non-coding region. We visualized the position of CpG sites relative to beginning of the first exon. The figure was created with BioRender.com **b,** Methylation value and correlation of 11 CpG site of the EDARADD gene. **c,** Correlation of the EDARADD promoter site (cg09809672) with age in the LiuY dataset **d,** Correlation of the EDARADD promoter site (cg09809672) with age in the training set of the Hannum clock and the largest blood training dataset of the Horvath multi-tissue clock (referred as Horvath 2012 dataset). **e,** Correlation of the EDARADD promoter site (cg09809672) with age in saliva of 42 twin pairs and in skin samples of the SkinBloodClock.

EDARADD had been used by microarray-based aging clocks, the CpG site cg09809672 annotated as promoter (TSS1500,5’UTR) in the arrays presented in the multi-tissue Horvath Clock, Hannum Clock based on blood, SkinBloodClock based on skin and blood cells and PhenoAge blood-based clock^6–9^. In the training set of Hannum Clock we found that the methylation level of the CpG site cg09809672 had a strong negative correlation with age (R: - 0.59, n=656). In the PhenoAge training set, this CpG site also had strong negative correlation (R: -0.64 n=456). The vast majority of the training samples in the multi-tissue Horvath clock were from blood and in the largest blood cohort of its training set we found that the promoter of the EDARADD gene had a strong negative correlation with age (R: -0.7, n=715, see Fig. 3d). We found that the cg09809672 CpG site of EDARADD was among the five common CpG sites of the examined four blood-based or blood-dominated methylation aging clocks. Intriguingly cg09809672 CpG site of EDARADD was in the same position as one CpG site in our multi-tissue aging clocks (see CpG 2 in Fig 3a.). We identified the cg09809672 CpG site in the Bhak et al. RRBS dataset in -1247 nt upstream from the first exon according to Havana/Ensemble transcript and it is also annotated TSS1500 in the arrays, while in MANE transcript we found it -62 nt upstream from the TSS in the 5’UTR as annotated in the array. Interestingly, it had the highest correlation among multi-omics aging clocks sites of EDARADD (Fig. 3b). To investigate this CpG site we tested 689 blood samples (referred to as Liu Y dataset), and we found that this CpG site is negatively correlated with age (Fig. 3c). The EDARADD promoter CpG site also negatively correlated with age in the training set of the Hannum clock and the largest blood training dataset of the Horvath multi-tissue clock (Fig. 3d). We observed the same in saliva of 42 twin pairs containing white blood cells with buccal epithelial cells^38^ (Fig. 3e). Analyzing 108 skin samples from 18-78 old individuals that were in the training set of SkinBloodClock, we found that this CpG site correlates with age in skin samples, and the SkinBloodClock model used it as a common predictor feature in skin and blood (Fig. 3e).

### Blood transcriptomic age acceleration and blood epigenetic age acceleration did not correlate

Using the multi-omics dataset, we measured both the epigenetics and transcriptomics age acceleration of individuals separately. Epigenetics and transcriptomics aging clocks were separately trained on 182 individual epigenomes and transcriptomes to predict age. We found a strong correlation between chronological age and the predicted age for both omics (Fig. 4ab). However, surprisingly, transcriptomics age acceleration and epigenetics age acceleration of the same individuals did not correlate significantly (Fig. 4cd). An explanation for this can be that the age prediction models may capture different aspects of aging.

**Fig. 4.**
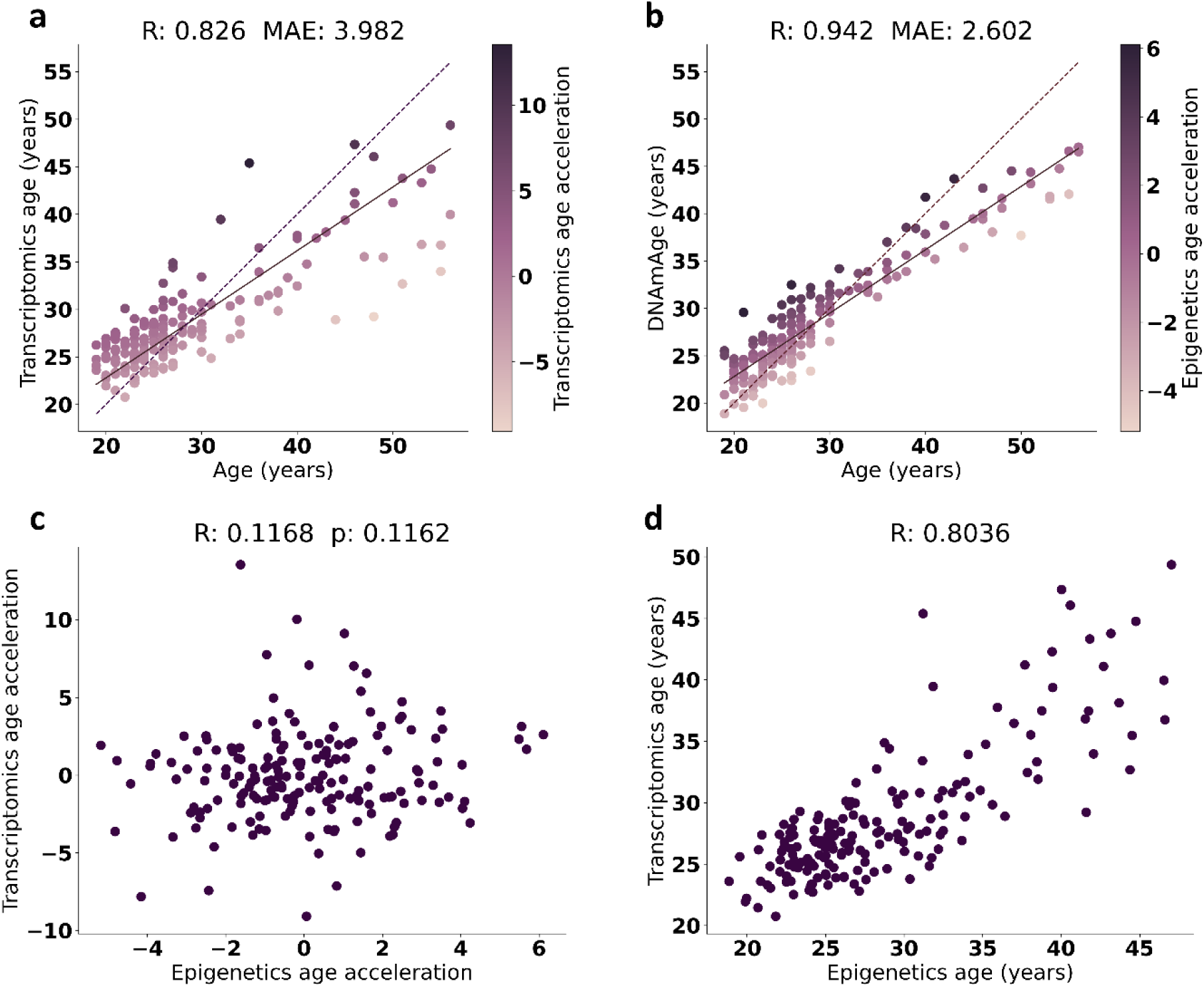
Age acceleration based on blood transcriptomes and epigenomes of the same individuals did not correlate. **a,** Age prediction performance of the transcriptomic aging clocks by using cross-validation. **b,** Age prediction performance of the epigenetics clocks by using cross-validation. **c,** Correlation analysis of the transcriptomics age acceleration and the epigenetic age acceleration of the same individuals. **d,** Correlation analysis of the transcriptomics age and epigenetics age of the same individuals.

## Discussion

The multi-omics clock showed that the adaptor gene EDARADD associated with the EDA-A1/EDAR/EDARADD Nf-kB canonical signaling pathway has a unique aging signature in both omics since not just the gene expression but the methylation also correlates with age, and we found hypomethylated promoter CpG sites. The first epigenetics age predictor using white blood cells from saliva used one promoter of EDARADD with a strong negative correlation (R = -0.8), and they found that the promoter is linear with age over a range of five decades and with NPTX2 they explained 73% of the variance in age in both sex^38^. The EDARADD contained one promoter of the identified 8 CpG sites that were independent of the cell-type composition of blood and one of the most informative sites with negative correlation on a training set consisting of 390 healthy blood samples^39^. There are microarray-based epigenetics clocks that use one CpG site of EDARADD for age prediction, namely the Horvath clock, the Hannum clock, the SkinBloodClock, and PhenoAge with a high negative correlation^6–9^. Genome-wide association study (GWAS) found that the EDARADD gene had the highest number of SNPs among the genes of different epigenetics clocks, they found 516 SNPs on EDARADD, and it’s associated with accelerated epigenetics aging^40^. Another study found that the promoter CpG site of EDARADD was hypomethylated in aged samples but in embryonic stem cells, and iPSCs it was highly methylated^41^. Methylation of that promoter site of EDARADD was significantly higher in newborns compared to adults and the elderly^42^. Hypomethylation of EDARADD is most significant in young ages and follows a linear progression from the very first years continuously through to older ages suggesting its role in early ages and development^43^. In forensic research using pyrosequencing, EDARADD hypomethylation has been widely used to estimate the age of individuals from blood, saliva, bone, and from teeth, nail, buccal swab^44–51^. EDARADD hypomethylation has been associated with Braak stage in prefrontal cortex of Alzheimer’s disease patients^52^. EDA/EDAR/EDARADD mediated Nf-κB signaling are associated with ectodermal development and impaired Nf-κB signaling in early development has a role in Ectodermal Dysplasia with characteristic of absence or abnormal ectodermal tissues for example dry, thin and wrinkled skin, loss of sweat glands and salivary glands, absent hair, teeth, and nails. Moreover, all inherited mutant EDARADD gene variants caused a reduction in the activation of EDA/EDAR/EDARADD Nf-κB, affecting ectodermal development^53–58^. Very recently, it has been shown that mutant EDA2R repressed Nf-κB in embryonic development suggesting its role in development^59^. EDA/EDAR/EDARADD pathway has a role in ocular surface development and its homeostasis^60^. It is shown that Gata6 developmental gene enhance EDARADD expression and activate NF-κB signaling in vivo, which in partly protect the hair follicle progenitor cells from DNA damage and premature apoptosis. In cultured keratinocytes, EDARADD rescues DNA damage, cell survival, and proliferation of Gata6 knockout cells. They observed a compensatory effect on the activity of NF-κB in vitro, cells were more dependent on this pathway in the stressful cell culture environment, and it was suggested that developmentally regulated transcription factors protect from DNA damage^61^. Others find that induced downregulation of EDAR, EDARADD, inhibited the activation of NF-κB pathway, resulting in impaired proliferation and increased apoptosis of keratinocytes^62^ EDAR overexpression promoted skin repair, improved wrinkles, and collagen deposition in skin and they observed the activation of wound healing genes. Moreover, they found EDAR expression was enhanced by Retinol, a widely used anti-wrinkle cosmetic^63^. Another study found that EDAR signaling contributed to human and murine skin repair further supporting the view that adult skin can reactivate embryonic processes under specific circumstances, and developmental pathways are reused and maintained in adult tissue as homeostatic control, for example in hair cycling and wound healing^64–66^. Another study showed that activation of certain embryonic development genes delayed cellular senescence and rejuvenated adult muscle progenitors without reprogramming their lineage specific fates or induce tumor^67^.

On the other hand, cancerous cells widely use EDARADD, which was overexpressed and as an oncogene promoted cell proliferation and activated Nf-κB signaling in colon cancer^68^. The expression level of EDARADD in TSCC tissues was markedly higher than in normal adjacent tissues and during in vitro knockdown of EDARADD induced apoptosis in TSCC cells and reduced the protein expression of NF-κBp65 and c-MYC^69^ which is a transcriptional regulator and mostly associated with proto-oncogenes activity and in induced pluripotent stem cell (iPSC) reprogramming as a Yamanaka factor^70^. EDARADD expression was significantly higher in high-grade ovarian tumors relative to normal tissue, EDARADD expression correlated with progression-free survival in patients with ovarian cancer and they hypotize EDARADD may be relevant to pathways underlying ovarian cancer initiation or progression^71^. EDARADD was significantly hypomethylated in cancer tissues in comparison to normal oral mucosa^72^. Other cancer studies have shown while analyzing tumor microenvironment, in liver cancer associated fibroblast EDARADD showed high expression and hypomethylation relative to non-cancer associated fibroblast. Hypomethylation of EDARADD correlated with worst survival in kidney, glioma, melanoma and in liver carcinoma^73^. Others found, that EDARADD was hypomethylated in normal prostate tissues and non-malignant prostate fibroblasts, and its correlated with age, while in cancer-associated fibroblasts it was hypomethylated and overexpressed, it was not correlated with age but it correlated with the survival , moreover tumors with low EDARADD DNA methylation and high EDARADD mRNA expression were consistently associated with shorter recurrence free survival., and the same was observed in lung cancer-associated fibroblasts, where only EDARADD correlated with worse progression-free survival among genes with aberrant promoter methylation, considered as a stromal biomarker, prognostic factor in non-small cell lung cancer^74,75^.

Multi-omics aging clock captured EDA and EDA2R as an age predictor feature and we found a positive correlation between the expression of these ligand and receptor pair with age in whole blood and PBMCs. Proteomics clock trained on circulating plasma proteins identified EDA2R as an age prediction feature in the proteomics level^76^. A very recent study showed that EDA2R is a ubiquitous hallmark of aging since its expression uniquely correlates with age in multiple solid tissues. Using more than 5000 whole blood samples from gene expression arrays, they found that EDA2R,EDA correlates with age and there was a positive association between the expression of EDA2R and C-reactive protein in the blood, and they suggest that both the elevated levels of EDA and EDA2R could lead to the activation of EDA-A2/EDA2R non-canonical Nf-κB signaling and it has a potential to trigger pro-inflammatory responses. In whole blood, they found both EDA-A1 and EDA-A2 mRNA but there were more EDA-A2 mRNA isoforms. They suggest that although blood-produced EDA may represent only a small fraction of total circulating EDA, however, they think aging-related coexisting conditions could further amplify EDA-A2/EDA2R pathway activation by increasing EDA ligand level^77^.

In summary, EDARADD is a widely recognized age predictor in different tissues, and its most acknowledged function is in EDA/EDAR/EDARADD mediated Nf-kB signaling, however, its gene regulatory activity is mostly associated with development, tissue homeostasis, and wound healing, furthermore in cancerous cells have been shown to overexpress EDARADD and activated Nf-κB

In aging research, understanding DNA methylation is essential, and differentiating stochastic variation from programmed is the first step to understanding regulatory mechanisms activated as we age. We found four age-related CpG sites near to the first exon in the promoter regions and other age-related CpG sites which were accumulated in the first intron. It has been shown that methylation of the first intron has inverse correlation with gene expression regardless of tissue and species^78^. We hypothesize that the EDARADD could be under epigenetics regulation during aging, and it might be associated with the activation of Nf-κB signaling, but its role in aging and age-related phenotypes is not known.

It is important to emphasize that we found methylation and gene expression alterations and we highlighted the importance of this signaling, however further investigations are needed to draw conclusions and understand its role in gene regulation, which might affect aging and rejuvenation.

A limitation of our study is that in the training dataset (Bhak et al. dataset), the oldest healthy participant was 40 years old, and every older individual exhibited adverse health phenotypes (suicide attempt or mild depressive disorder)^34^. The Bhak et al. study created classifiers that distinguished healthy individuals from depressed ones. We found no overlap between the features of these classifiers and the features of the multi-omics aging clocks (Supplementary Table 7). Also, we found that a large proportion of the 32 genes of the multi-omics aging clocks genes highly correlated with age in the healthy PBMC samples of the AIDA and the Morandini et al. datasets.

Another limitation of our study is that we did not validate the multi-omics aging clocks in an external dataset since we were unable to find a suitable publicly available blood multi-omics dataset with age and RRBS data that has sufficient genome coverage to obtain age-related methylation information.

In this study, we developed age prediction models trained on multi-omics (epigenetics and transcriptomics) data from the peripheral blood of 182 individuals, and analyzing them, we concluded that multi-omics clocks are useful tools for investigating aging and biological age at the multi-omics level.

## Methods

### Datasets and pre-processing

We processed the raw bisulfite sequences of the Bhak et al. study^34^(PRJNA531784) as we described earlier^16^. Shortly, bisulfite-sequencing raw fastaq files of the 182 peripheral blood samples were downloaded from NCBI SRA (PRJNA531784). The samples were collected from 56 suicide attempters, 39 patients with major depressive disorders and 87 healthy controls. Trimming and quality control were made by Trim Galore. For the methylation calling we used Bismark with Bowtie 2 mapping to the GRCh38 human genome assembly^79^.

We also acquired the RNA-seq files of the same 182 samples (PRJNA531784). We trimmed the reads with Trimmomatic and used HISAT2 to align them to the GRCh38 human genome assembly. The gene expression matrix was created by counting the uniquely mapped reads in each gene using featureCount with the Ensembl GRCh38.p14 human gene annotation. To take sequencing depth into account, the raw gene expression matrix was normalized by dividing each count by the total number of aligned reads in each sample.

For the pseudobulk samples, we used the AIDA Data Freeze v2 dataset^35^ (https://cellxgene.cziscience.com/collections/ced320a1-29f3-47c1-a735-513c7084d508). We used the normalized gene expression matrix of bulk PBMC of the Morandini et al. dataset^36^ that is available in the study as a supplementary table. Furthermore, we used the training set of the Hannum clock (GSE40279), the largest blood training dataset of the Horvath multi-tissue clock (GSE41037), the blood cohort from Liu et al. (GSE42861), blood samples of 42 twin pairs (GSE28746), and the training set of the SkinBloodClock (E-MTAB-4385).

### Training of the multi-omics aging and transcriptomics clocks

The chronological age prediction models (bulk multi-omics and transcriptomics clock) were trained by using the Python implementation of GLMNet which automatically fine-tunes the lambda parameter. The best value for the alpha parameter that chose gene expression features was alpha=0.01. In the separately trained transcriptomics clock (with172 genes) the best alpha was 0.1. To prevent overfitting, we used leave-one-out cross-validation. We analyzed the feature sets of these models in the analysis. The multi-omics models selected 4150 methylation features and 33 gene expression features in total. For the further analysis we dropped one of the gene expression features which was not annotated (ENSG00000287593, “novel transcript”).

### Creating pseudobulk samples

We used AIDA dataset, in which there are 1,265,624 circulating immune cells, and 32 cell types from 625 healthy donors with ages between 19 and 75 years. We created pseudobulk samples by aggregating raw counts of cells per donor. To avoid the batch effect caused by differences in the number of cells among individuals we divided each count by the total number of reads in each sample. For the figures, we multiplied the values by one million and we used Counts per Million (CPM) to calculate the correlation between the gene expression and chronological age.

### Age acceleration

The age accelerations were calculated as the residual from regressing predicted omics age on chronological age for each omics separately, resulting in „epigenetics age acceleration” and „transcriptomics age acceleration”.

### Overrepresentation analysis

Overrepresentation analysis (ORA) is a widely used analysis for interpreting gene lists of interest. Mostly it is used with differential gene expression analysis (DEG) where the gene list is the DEG genes. In this study, we used this method to extract biological information from the features of the machine learning models that capture biomarkers that correlate with age. Using this gene set we performed ORA in g:Profiler (https://biit.cs.ut.ee/gprofiler/gost). g:Profiler^80^ created the functional profiling of our gene list with the following setting: background genes were corrected for the genes in the training set. Searching was performed in all GO Molecular Functions, Reactome, KEGG and WikiPathways. We excluded results supported by electronic GO annotations. For multiple samplings, we used Benjamini-Hochberg correction.

Cytoscape with Enrichmap was used to determine the network of the overrepresented pathways. Clustering and annotation were performed by the default algorithm of Cytoscape^81^.

### Promoter annotation

The transcription start site determined by the first nucleotide of the first exon, start of the coding region in the transcripts. Promoter region on the gene was determined as [-1500, +500] base pair from transcription start site following the direction of transcription. We used the MANE (Matched Annotation from NCBI and EMBL-EBI) Project^37^ transcripts matched GRCh38 and are 100% identical between Ensembl and Refseq, GENECODE for 5’ UTR, CDS, splicing and 3’ UTR and supported by experimental data, transcription start site is supported by CAGE sequencing. For further investigation, we used another transcript, from EMBL-EBI, and it was annotated by Havana and Ensembl automatic annotation.

### Data and code availability

The training dataset and the code will be made publicly available upon publication. We selected a representative multi-omics aging clock model from the 182 models trained on the Bhak et al. dataset via leave-one-out-cross-validation (see above). The model coefficients of the multi-omics aging clock are available in Supplementary Table 5.

## Author contributions

C.K. conceived the study and supervised the work. B.V. retrieved data, developed the multi-omics (and other) aging clocks and performed the data analysis, discussed the results. Both authors interpreted the results, and wrote the manuscript.

## Acknowledgments

The project was supported by the HUN-REN (TKCS-2024/37), and European Union project RRF-2.3.1-21-2022-00004 within the framework of the Artificial Intelligence National Laboratory. The project was also supported by the National Research, Development and Innovation Office – NKFIH, FK-146113.

